# The dimensionality of neural coding for cognitive control is gradually transformed within the lateral prefrontal cortex

**DOI:** 10.1101/2024.02.05.578918

**Authors:** Rocco Chiou, John Duncan, Elizabeth Jefferies, Matthew A. Lambon Ralph

## Abstract

Implementing cognitive control relies on neural representations that are inherently high-dimensional and distribute across multiple subregions in the prefrontal cortex (PFC). Traditional approaches tackle prefrontal representations by reducing them into a unidimensional measure (univariate amplitude) or using them to distinguish a limited number of alternatives (pattern classification). By contrast, representational similarity analysis (RSA) enables flexibly formulating various hypotheses about informational contents underlying the neural codes, explicitly comparing hypotheses, and examining the representational alignment between brain regions. Here, we used a multifaceted paradigm wherein the difficulty of cognitive control was manipulated separately for five cognitive tasks. We used RSA to unveil representational contents, measure the representational alignment between regions, and quantify representational generality *vs.* specificity. We found a graded transition in the lateral PFC: The dorsocaudal PFC was tuned to the information about behavioural effort, preferentially connected with the parietal cortex, and representationally generalisable across domains. The ventrorostral PFC was tuned to the abstract structure of tasks, preferentially connected with the temporal cortex, and representationally specific. The middle PFC (interposed between dorsocaudal and ventrorostral PFC) was tuned to individual task-sets, ranked in the middle in terms of connectivity and generalisability. Furthermore, whether a region was dimensionally rich or thin co-varied with its functional profile: Low dimensionality (only gist) in the dorsocaudal PFC dovetailed with better generality, whereas high dimensionality (gist plus details) in the ventrorostral PFC corresponded with better ability to encode subtleties. Our findings, collectively, demonstrate how cognitive control is decomposed into distinct facets that transition steadily along prefrontal subregions.

**Significance:** Cognitive control is known to be a high-dimensional construct, implemented along the dorsocaudal-ventrorostral subregions of PFC. However, it remains unclear how prefrontal representations could be dissected in a multivariate fashion to reveal (1) what information is encoded in each subregion, (2) whether information systematically transforms across contiguous PFC subregions as a gradient, (3) how this transformation is affected by functional connectivity. Here we shed light on these issues by using RSA to decode informational composition in the PFC while using participant-specific localisers to facilitate individually-tailored precision. Our findings elucidate the functional organisation of PFC by revealing how a trade-off between dimensionality and generalisability unfolds in the PFC and highlight the strength of RSA in deciphering the coding of cognitive control.

## Introduction

The prefrontal cortex (PFC) is an anatomically complex and functionally heterogeneous brain region. Anatomically, the PFC contains multiple subregions, each receiving input and sending output to a unique set of other areas (e.g., Thiebaut de Schotten et al., 2017; Jung et al., 2022). Perhaps as a result of varied connectomic fingerprints, the PFC is functionally manifold – some subregions are ubiquitously involved in diverse situations regardless of stimuli and operations (Duncan, 2010) while others exhibit proclivity for certain types of stimuli/tasks (e.g., Goldman-Rakic, 1996). For decades, scientists have endeavoured to understand how PFC subregions are systematically organised so as to identify an organisational axis that best ascribes functions to structures. One of the prominent findings is that functional profiles gradually vary along the dorsocaudal-ventrorostral axis of PFC (for review, see Badre and Nee, 2018; Constantinidis and Qi, 2018; Abdallah et al., 2022). At one end of the axis, the dorsocaudal PFC preferentially encodes concrete/perceptible representations to prepare for action (e.g., mapping a visible stimulus location to finger reaction). At the other end, it has been proposed that the ventrorostral PFC preferentially encodes abstract representations (entities not directly perceivable, such as semantics or abstract relationship among stimuli). Critically, regions situated between dorsocaudal and ventrorostral PFC vary in a stepwise manner, with those midway between the extremes (e.g., the middle frontal gyrus) preferentially representing the current focus of attention or the goal of behaviour (Woolgar et al., 2011). These middle frontal regions might integrate information from dorsocaudal and ventrorostral sources (Nee and D’Esposito, 2016).

While the dorsocaudal-ventrorostral axis has offered an anchoring framework for mapping out the relationship between functions and subregions, it is nevertheless incomplete. Below we discuss the insufficiency of existing research and how the present study was performed to address these problems. ***First***, a lack of crosstalk between literatures: Traditionally, research on executive functions tends to employ highly abstract rules and stimuli (such as arbitrarily mapping visual shapes to motor outputs) and consider semantic meaning as ‘contamination.’ However, a substantial swath of the PFC is also involved in semantic tasks, especially during goal-directed retrieval/manipulation of semantics (e.g., Chiou et al., 2018; Gao et al., 2021). To fully understand the correspondence between PFC structures and functions, it is necessary to incorporate both semantic and non-semantic components in a single experimental design. ***Second***, the limitation of univariate analysis: As discussed by Freund *et al*. (2021b), the majority of neuroimaging studies on cognitive control employed univariate contrasts, which inherently condenses potentially multifaceted processes into a uni-dimensional measure (activity amplitude) and therefore loses information. Contrary to univariate analysis, representational similarly analysis (RSA) has the ability to decompose the informational contents distributed across a region’s neuronal population by revealing the dimensions that define this representation. Using RSA, we can relate a region’s functionality (domain-general *vs.* -specific) to its dimensionality (limited *vs.* multifaceted). ***Third***, a lack of precision when defining regions of interest (ROI): As discussed by Fedorenko (2021), the topography of task-induced activation can be idiosyncratic across individuals. As a result, an analysis relying on anatomical templates or group-level clusters loses sensitivity and can result in inaccurate estimates for an individual. To combat this imprecision issue, it has been suggested that using a functional localiser to individually identify ROIs greatly improves accuracy (Shashidhara et al., 2020). In the present study, we tackled these issues by manipulating various semantic and non-semantic variables to create a design conducive for conducting RSA; a localiser task was employed to precisely demarcate individual-specific prefrontal ROIs.

Our RSA decoding reveals an informational gradient in the PFC, transitioning from representing cognitive effort, via individuality of task-sets, to semantic meaning. Different subregions along the gradient preferentially exchange information with distinct regions outside the PFC and have different representational dimensionalities that dovetail with their functional properties (general *vs.* specific; concrete/motoric *vs.* abstract/conceptual). Together, these results reveal an organisational principle that specifies *how* and *where* information transforms within and beyond the PFC.

## Materials and Methods

### Participants

Twenty-five volunteers gave informed consent before the scanning session. The sample consisted of 15 females and 10 males, with an average age = 28 years-old and SD = 6. All volunteers spoke English as their native language and were right-handed. All participants completed a screening questionnaire about magnetic resonance imaging safety before the scanning session began. None of them reported having any previous brain injury, and not having any neurological or psychiatric condition. This study was reviewed and approved by the Northwest Research Ethics Committee.

### Design and stimuli

After the structural scan, volunteers completed eleven runs of echo-planar imaging in a single session. Run 1 was the localiser experiment and its associated results have been published in Chiou *et al*. (2022). This localiser was designed to probe cognitive control for semantic and perceptual processes, and its individual-level contrast results were used to localise control-related ROIs for each participant. Run 2 to 11 was the RSA decoding experiment. As the primary goal of our study was to compare the similarities and differences among prefrontal subareas, decoding analysis on the data of Run 2 to 11 was primarily focused on the individual-specific prefrontal ROIs defined by the localiser contrasts.

Details about the localiser task are comprehensively described in Chiou *et al*. (2022). Below we report the key information: We used a 2-by-2 factorial design in which different types of processing (semantic processing on lexical stimuli *vs.* visuospatial processing on pictorial stimuli) and the extent of difficulty required to achieve a correct response (easy *vs.* hard) was orthogonally manipulated. There were four items displayed in each trial (either four English words or four geometric patterns), and one of the four items served as an ‘oddball’, either semantically incongruous with other words (belonging to a separate semantic category from the rest) or perceptually incongruous with the others (having a distinct visual configuration). Difficulty level was manipulated by varying the closeness of semantic categories or the similarity of visual patterns. Participants had to indicate the oddball with a button response. The localiser experiment had a block design, with each of the four conditions having 12 blocks; each block consisted of three trials; each trial started with a fixation cross (0.1 sec), followed by a quadruplet of stimuli (3.9 sec). Together with inter-block intervals, the localiser task had a total duration of 647.5 sec. The experiment was programmed using E-Prime 2.0 (2013, Psychology Software Tools). When performing the task, participants responded by pressing one of the four designated buttons on an MR-compatible response-pad using their right fingers. All text stimuli were white colour, displayed on a black background; text was Arial typeface, 24-point in font size. Stimuli were displayed using an MRI-specialised LCD screen (1,920 × 1,080; NordicNeuroLab) and projected onto a coil-mounted device for viewing stimuli.

The RSA decoding experiment contained 10 runs of scanning (Run 2 to 11). We employed a 5-by-2 factorial design that fully crossed different types of task processing (five tasks: Semantic Association, Synonym Decision, Colour Knowledge, Numerical Comparison, and Shape Matching) with the difficulty with which to attain a correct answer (easy *vs.* hard). The overall structure of the experiment is illustrated in Figure 1A. As the Figure shows, in each trial there was a reference item on the top (a word, a number, or a shape) and three options beneath. In different tasks, participants were asked to engage in different types of cognitive operations and choose an answer in relation to the reference. The specifics of task requirements were: (***1***) For the Semantic Association task, participants evaluated the relationship between the reference noun word and three options, and chose the option (target) semantically most associated with the reference. We manipulated the strength of association between the easy and hard condition of this task such that semantic relationship was obvious and straightforward in the easy condition (e.g., *salt* and *pepper*), but ambiguous and indirect in the hard condition (e.g., *salt* and *grain*). To ensure the adequacy of stimuli, eight additional volunteers (none took part in the MRI study) were recruited to rate semantic association between the reference and options on a 5-point scale and the data showed that, as intended, the targets were significantly more strongly associated with the references in the easy context (mean±SD: 4.56±0.27) than in the hard condition (3.39±0.65); the difference was found at the individual level (by-item analysis: *p* < 10^-9^ for all raters) and the group level (by-subject analysis: *p* < 0.001), and the non-target options (foils) were significantly less associated with the references than targets (*p*s < 0.001). We also controlled for word length (by letter count) and lexical frequency (using the corpus of Van Heuven *et al*., 2014) for all words used in the task so that ‘easy *vs.* hard’ did not differ on these linguistic variables. (***2***) For the Synonym Decision task, participants compared the conceptual similarity between the reference noun and the three options, and chose the target most synonymous with the reference. We manipulated the ease with which to access lexico-semantics by using concrete nouns (tangible/perceptible concepts, such as *box* or *kettle*) in the easy condition and abstract nouns (e.g., *onus* or *deceit*) in the hard condition, based on the well-documented findings that lexical access to abstract words is cognitively more taxing than to concrete words (e.g., Hoffman et al., 2015). Based on the concreteness ratings from Brysbaert *et al*. (2014), we ensured that the concrete words used in the easy condition were rated as more concrete than the abstract words in the hard condition (*p* < 10^-25^) while there was no difference in word length and lexical frequency (both *p*s > 0.45); we recruited eight additional participants to rate the synonymy of words and ensured that the targets and references were judged as more synonymous with each other compared to foils (*p*s < 0.001 for both concrete and abstract words). (***3***) For the Colour Knowledge task, participants needed to retrieve the knowledge about canonical colour of objects; the references were nouns associated with a typical colour attribute (e.g., *banana* is typically yellow), and the task was to choose the option that shared a similar colour to the reference. The difficulty manipulation here concerned the relationship between the references and foils – in the hard condition, the foils were semantically highly related to the references, which made them highly distracting as participants must actively suppress a tendency to respond to those semantically related (yet task-irrelevant) foils and concentrate on identifying the semantically unrelated targets using the arbitrary task rule (i.e., pairing words using colour similarity) (for precedent of this task design, see Badre et al., 2005). In the easy condition, by contrast, the targets and foils were all semantically unrelated to the references (despite the targets and references sharing similar colours), hence reducing the interference of foils. We collected eight additional volunteers’ ratings to ascertain that the foils were significantly more related to the references in the hard than easy condition (*p*s < 0.0001) while the targets and references were minimally related despite sharing similar colours. Word length and lexical frequency were controlled for to ensure these factors did not confound the key manipulation (both *p*s > 0.27). (***4***) For the Numerical Comparison task, volunteers compared the numerical distance between the reference number and three options and identified the option numerically nearest to the reference. The numbers were 3-digit numerals, ranging from 200 to 999. In the easy condition, the target had an obviously closer distance to the reference compared to foils, whereas in the hard condition, the distance was not obvious, which necessitated careful comparison to identify the target. (***5***) For the Shape Matching task, volunteers viewed simple geometric shapes and were asked to compare their perceptual sizes. In the easy condition, it was easy to identify the target that had a perceptually very comparable shape to the reference, whereas in the hard condition careful visual inspection was necessary to select the target because all options had a similar size to the reference. For the three conditions in which lexical stimuli were used (i.e., Semantic Association, Synonym Decision, and Colour Knowledge), we carefully cherry-picked 1,200 words as the references, targets, and foils to ensure that (*i*) each word only appeared once in the entire experiment and (*ii*) none of the words was repeatedly used to avoid the effect of residual memory from previous trials interfering between conditions/tasks. In all tasks, the targets were equally likely to appear in any of the three locations, and thus equally likely to be reacted to using any of the three buttons.

**Figure 1.**
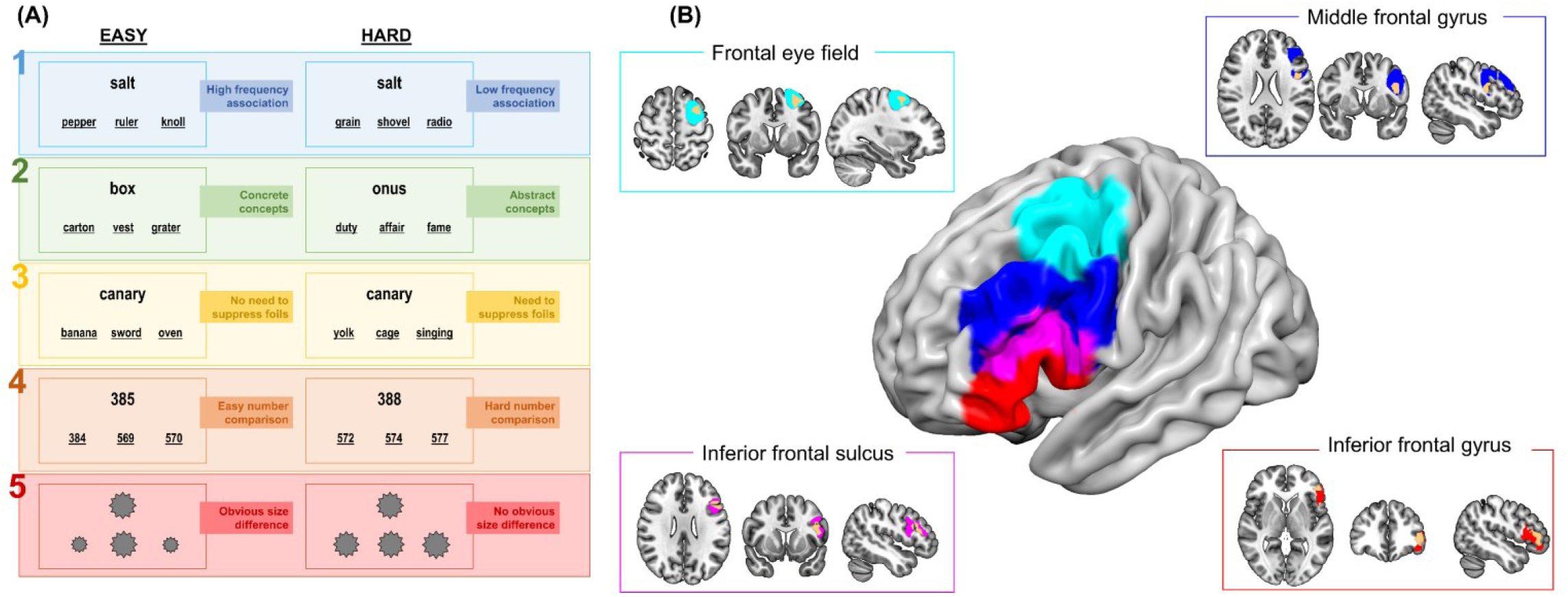
(**A**) The design of the RSA decoding experiment contained ten task-conditions. Specifically, there were five tasks (1: Semantic Association; 2: Synonym Decision; 3: Colour Knowledge; 4: Numerical Comparison; 5: Shape Matching) and for each task we manipulated the level of difficulty to achieve a correct response (Easy vs. Hard), yielding ten conditions in total. (**B**) Selection of voxels for defining individual-specific ROIs had the following steps: *First*, functionally defined group-level clusters from Fedorenko *et al*. (2013) and Jackson (2021) were used to constrain the scope of selection and ensure consistency with prior literature. *Second*, based on the functional clusters’ alignment with gyri and sulci structures, the clusters were divided into four sections: frontal eye field (FEF; cyan); middle frontal gyrus (MFG; blue), inferior frontal sulcus (IFS; magenta), and inferior frontal gyrus (IFG; red). *Third*, using the results of localiser experiment, for each individual participant we selected the top 200 voxels, typically contiguous, most responsive to the ROI-defining contrast within each of the four PFC divisions. Examples results from one participant are illustrated as the orange patches in the inset boxes.

Each run of scanning consisted of 20 blocks of trials; prior to the onset each block, an instruction was presented for 3 sec to inform the upcoming task (i.e., Semantic Association, Synonym Decision, Colour Knowledge, Numerical Comparison, and Shape Matching). Each block consisted of 5 trials; each trial began with a fixation dot (0.4 sec), followed by a quadruplet of stimuli (one reference and three options) displayed for 3.5 sec, yielding 19.5-sec duration for each block of trials, and each block was succeeded by a 2-sec interval before the onset of next instruction. We counterbalanced the order in which the 10 conditions appeared across runs and participants so that each condition was equally probable to appear in any slot of the sequences, and equally probable to be preceded and succeeded by any other conditions, with each condition having 20 blocks of trials in total and each run lasting 489 sec. The order of stimuli within each condition were randomised. Participants were instructed to respond as quickly and accurately as possible within a trial’s time-limit. All stimuli were (words, numbers, and shapes) were presented in white colour against a black background. Identical devices and parameters for stimuli display and response collection to the localiser experiment were used in the present RSA decoding experiment. It is worth noting that our experimental design, using a fully counterbalanced factorial structure, ensured that all conditions were equally likely to appear in any run of scans, with each condition having same number of blocks, and evenly distributed across the timeslots within a run, which prevented some confounding biases in RSA that an unbalanced design might have (for discussion, see Freund et al., 2021a).

### Regions of interest

A common practice of multivoxel pattern analysis (MVPA) is to investigate distributed patterns of activity within an ROI using anatomical templates (e.g., Woolgar et al., 2011) or atlas parcellations (e.g., Freund et al., 2021a). Although this method offers high consistency across individuals (i.e., exactly the same voxels are selected for all participants), it omits the heterogeneity of brain activities between individuals. As demonstrated in recent reports (Shashidhara et al., 2020; Fedorenko, 2021), the loci of functional responses are somewhat idiosyncratic for each person; the conventional method of atlas parcel provides a ballpark estimate but is not sensitive enough to capture subtle idiosyncrasy, which has been suggested to be particularly problematic for decoding prefrontal activation (Bhandari et al., 2018). To tackle this issue, in the present study, when defining the ROIs we employed the ‘group-constrained yet individual-specific’ method advocated by Fedorenko (2021), which has been demonstrated to successfully capture individual variation while ensuring all individual loci are situated within a predefined target realm (e.g., Fedorenko et al., 2013; Blank and Fedorenko, 2017; Mineroff et al., 2018). The steps of defining ROIs are: (***i***) First, we specified the scope for selecting voxels using the functional masks from Fedorenko *et al*. (2013) and Jackson (2021). Fedorenko *et al*. (2013) focused on *domain-general* executive control and identified a group of prefrontal regions, mostly in the dorsolateral prefrontal cortex (DLPFC) that encompasses the frontal eye field (FEF) and middle frontal gyrus (MFG). Jackson (2021) conducted a meta-analysis surveying the literature of *domain-specific* semantic control (executive operations specifically on semantic representations) and identified several regions in the ventrolateral prefrontal cortex (VLPFC) that encompassed the entirety of the inferior frontal gyrus (IFG) and inferior frontal sulcus (IFS, a site superior to the IFG). It is worth noting that the two studies that we used here to define the boundaries of voxel selection are highly representative of the literature across a wide variety of paradigms (i.e., seven tasks probing domain-general control in 197 subjects, and 87 fMRI studies on diverse ways of semantic control). The masks are openly downloadable from the original studies. (***ii***) Next, using the macroanatomy of AAL atlas (Tzourio-Mazoyer et al., 2002), the clusters of domain-general control (Fedorenko *et al*., 2013) and domain-specific control (Jackson, 2021) were parcellated into four divisions based on the alignment with gyri/sulci structures (for illustration, see Figure 1B): (***1***) the FEF parcel contains exclusively domain-general voxels and is most dorsocaudal; (***2***) the MFG parcel contains exclusively domain-general voxels and is ventral/rostral to the FEF; (***3***) the IFS is the conjunction zone between the domain-general and domain-specific clusters and is located ventral/rostral to the MFG; (***4***) the IFG contains exclusively domain-specific voxels and is most ventrorostral. (***iii***) Finally, for each of the four prefrontal parcels and for every participant, individual localiser results were used to identify the top 200 voxels showing biggest contrast parameters (see Figure 1B for examples). Specifically, using the main effect contrast of ‘Hard > Easy’ (aggregating across levels of the semantic condition and visuospatial condition), we identified the 200 voxels showing largest response to the overall effect of difficulty. The selected voxels from each parcel were used to construct individual-specific ROIs for subsequent MVPA using the data from the RSA experiment. It is worth stressing that the number of voxels was predefined before we conducted any analysis; the number (200 voxels) was based on prior studies showing that increasing the size of an ROI up to 200 voxels robustly improved decoding before it reached a plateau as more voxels were added (Shashidhara et al., 2020). With this protocol to define ROIs, we were able to adapt to the heterogeneous distribution between individual results while considering prefrontal anatomy and ensuring consistency with prior literature. Moreover, given that differing number of voxels is known to be a confounding factor in MVPA (Todd et al., 2013), our approach avoids this problem by ensuring equivalent number of voxels in each ROI and thus creating a level playing field for comparing between ROIs.

For each individual, we additionally defined two ROIs in the posterior/superior parietal cortex (PPC) and the posterior middle temporal gyrus (pMTG) for the ‘representational alignment’ analysis to investigate how the four prefrontal ROIs share representational structures with remote areas and whether there is any systematic change along the dorsocaudal-ventrorostral axis. Whereas the PPC is part of the dorsal-attention network and heavily involved in visuospatial tasks (e.g., Yeo et al., 2011; Mengotti et al., 2020), the pMTG is part of the semantic-control system and frequently involved when semantic operations become more difficult (Davey et al., 2016; Jackson, 2021; Chiou et al., 2022). Our previous report of the localiser data (Chiou et al., 2022) also showed that the PPC and pMTG are, respectively, the regional peaks for visuospatial control (MNI: *-28*, *-58*, *52*) and semantic control (*-62*, *-42*, *-8*). Given that the PPC and pMTG have distinct functional profiles, we used different localiser contrasts to define the ROIs: (***1***) For the PPC, confined within the cortical mask of the superior parietal lobule (Harvard-Oxford Atlas: Desikan et al., 2006), we selected the top 200 voxels showing largest responses to the localiser contrast of ‘Visuospatial-Hard > Visuospatial-Easy’ for each participant to construct individually-tailored ROIs; (***2***) For the pMTG, confined in the posterior/temporo-occipital divisions of middle temporal gyrus, we individually selected 200 voxels showing largest responses to the localiser contrast of ‘Semantic-Hard > Semantic-Easy’. Akin to the way we defined prefrontal ROIs, our approach accommodated individual variation while ensuring the equivalent number of voxels in each ROI, and all chosen voxels were near the group-level average and situated in a predefined target territory.

In Extended Data (Figure 1–1), we report a univariate analysis on the localiser results, comparing the four prefrontal ROIs’ preferences for semantic *vs.* visuospatial processing. Results corroborate the previous literature on prefrontal functions (e.g., Nee and D’Esposito, 2016) – the dorsolateral PFC preferentially responds to visuospatial processing, while the ventrolateral PFC preferentially responds to semantic processing. This justified our labelling (in subsequent figures of our study) of the FEF (most dorsocaudal) as ‘visual-sensitive’, the IFG (most ventrorostral) as ‘semantic-sensitive’, and the MFG and IFS (situated in the intermediate territory) as ‘domain-general’ (owing to these two regions responding equally strongly for visuospatial and semantic processing).

### fMRI acquisition and pre-processing

The MRI protocol and pre-processing parameters are identical to those reported in Chiou *et al*. (2022). A Siemens 3-Tesla PRISMA system was used to acquire all of the MRI data. Anatomical images were attained with a T_1_-weighted Magnetisation Prepared Rapid Gradient Echo (MPRAGE) sequence [repetition time (TR) = 2250 ms; echo time (TE) = 3.02 ms; inversion time = 900 ms; 230 Hz per pixel; flip angle = 9°; field of view (FOV) 256 × 256 × 192 mm; GRAPPA acceleration Factor 2]. Task-related data were obtained with a multi-echoes multi-band (MEMB) blood oxygenation level dependent (BOLD)-weighted echo-planar imaging (EPI) pulse sequence. This MEMB sequence has the strengths that it acquired four functional volumes for each TR (multi-echoes, enabling capturing signals that peaked at early and late echo-times that are often overlooked by conventional protocols) and simultaneously recorded two slices during the acquisition of each volume (multi-band, expediting the acquisition rate). The parameters included: TR = 1,792 ms; TE_1_ = 13 ms, TE_2_ = 23.89 ms, TE_3_ = 34.78 ms, TE_4_ = 45.67 ms; flip angle = 75°; FOV = 192 mm × 192 mm, MB-Factor = 2, in-plane acceleration = 3. Each EPI volume consisted of 46 axial slices, acquired from the top and middle slice in descending order, with voxel size of 3 × 3× 3 mm and FOV of 240 × 240 × 138 mm. A series of 362 and 273 functional volumes per run, respectively, were acquired for the localiser experiment and the RSA decoding experiment.

All raw DICOM data were converted to NifTi format using dcm2niix. The T1 structural images were processed using the standard processing pipeline of the FSL package’s ‘fsl_anat’ function (Ver5.0.11; https://fsl.fmrib.ox.ac.uk/). This procedure involves the following six steps: (*i*) Reorient images to standard MNI space (‘fslreorient2std’), (*ii*) automatically crop image to remove the neck (‘robustfov’), (*iii*) bias-field correction to fix field inhomogeneity (‘fast’), (*iv*) registration into the MNI space (‘flirt’ and ‘fnirt’), (*v*) brain extraction (using ‘fnirt’ nonlinear method) and (*vi*) tissue-type segmentation to separate white-/grey-matter and other structures (‘fast’). Each T_1_-image was individually inspected for accuracy after being normalised into the MNI space. The functional EPI data were pre-processed using a combination of tools in FSL, AFNI (Ver18.3.03; https://afni.nimh.nih.gov/), and a specialised Python package to perform TE-dependent analysis (Kundu et al., 2012; Kundu et al., 2013; Kundu et al., 2017). Despiked (‘3dDespike’), slice-time corrected (‘3dTshift’, matched to the middle slice), and realigned (‘3dvolreg’) images were submitted to the “tedana” toolbox, which took the time-series data from all of the four TEs and decomposed the data into BOLD-signals and noises (non-BOLD). Specifically, decomposition was based on whether a signal series depended on the four echo-times – with the strength of multiple echo-times, the algorithm was able to tell apart noises that fluctuated randomly and independently of the timing of four TEs (e.g., scanner’s drift, cardiac/respiratory noises, head motion) from signals that systematically varied with the timing of TEs (e.g., real functional data of BOLD). Data of the four echo-times were optimally integrated, weighted based on the intensity of T2* signal in each voxel and separated from the TE-independent/ non-BLOD noises (Kundu et al., 2017). Finally, the optimally combined images were co-registered into individual’s T_1_ structural scan (using FSL’s ‘flirt’), normalised to the MNI space (using FSL’s ‘fnirt’ warps and ‘flirt’ transform), and smoothed with a 6 mm FHWM Gaussian kernel.

### Multivoxel pattern analysis (RSA and classification)

Prior to multivariate decoding, we used SPM12 to construct general linear models (GLM) to fit the data of the localiser experiment and the RSA experiment, convolving each participant’s design matrix with a canonical haemodynamic response function. For the localiser experiment, four task regressors were created in the GLM (Semantic-Easy, Semantic-Hard, Visuospatial-Easy, Visuospatial-Hard), which enabled the relevant contrasts for defining ROIs. For the RSA experiment, 10 task regressors were created to enable constructing a 10×10 representational distance matrix in RSA (Association-Easy, Association-Hard, Synonym-Easy, Synonym-Hard, Colour-Easy, Colour-Hard, Number-Easy, Number-Hard, Shape-Easy, Shape-Hard). Based on the established methods (Kriegeskorte et al., 2008; Nili et al., 2014), for the off-diagonal elements of the matrix, we calculated the neural similarity between each pair of experimental conditions as the Pearson correlation of the two conditions’ vectorised patterns of neural activities; the similarity results were converted into a distance measure (1 – Pearson’s *r*) to represent how distant the neural patterns are from one another. In addition to constructing matrices using the neural data, we also constructed various theoretical matrices based on different assumptions about the relationship between task conditions, and used them to account for the neural results. Details of the theoretical models are presented later in the Results. Based on the standard practice of RSA (e.g., Nili et al., 2014), we used a rank-based correlation index Kendall’s τ (τ-a) to measure the similarity between different matrices. We also computed Spearman’s *ρ* and found that it yielded entirely consistent results with τ-a; however, we chose to report τ-a as Nili *et al*. (2014) recommend that, compared to Spearman’s *ρ*, τ-a offers a more conservative and detailed prediction.

While τ-a has been widely used as a standard indicator when researchers compare hypotheses to see which ones better explains the neural representations, more stringent indices are needed to evaluate the unique variance explained by the hypothesis under consideration while explanatory influences from all other sources are removed. This concern is especially crucial in one of our analyses in which we sought to compare among the prefrontal subregions to investigate whether their neural coding was loaded with certain information. In this analysis, the neural matrices served as the candidates for adjudication, and the theoretical matrices are the targets of prediction – e.g., among the four regions, which one carried most information about reaction times, and which least? However, an important consideration here was that we had to control for the confounding influences from all other subregions when gauging the unique variance the target brain region could explain. Given that our four ROIs are spatially contiguous, it is critical to statistically rule out the contamination from other regions to get a clean estimate for each region. To this end, we used two different approaches: (***1***) We computed the partial correlation (Spearman’s *ρ*), which focused on the unique relationship between the neural and theoretical matrices (e.g., the similarity between IFG and a hypothesis model) while excluding the co-variance they shared with all other prefrontal areas (i.e., the FEF, MFG, IFS). (***2***) We used a multiple-regression approach, following the example of Freund *et al*. (2021b). For multiple regression, the off-diagonal elements of four candidate matrices (i.e., the neural matrices of four prefrontal ROIs) simultaneously entered the regression as the predictors. The neural data were warped into vectors, rank-transformed, and normalised into *z*-scores, while one of the theoretical matrix served as the dependent variable. With this approach, the regression coefficient *β* of each regressor represented the explanatory power unique to each regressor while the effects from all other regressors were removed. It is worth noting that RSA was performed at the individual level to ascertain that fine-grained, voxel-wise representations were preserved for every person, and group-level averaging and statistical tests (e.g., comparing the sizes of correlations or regression coefficients) were performed only after representational distance matrices were computed for each participant and for each ROI. Finally, in addition to relating neural matrices to theoretically-constructed matrices (which is a more common practice in RSA), we also performed ‘representational alignment’ analyses, which quantified the extent to which brain regions’ representational structures aligned with one another (cf. Ito and Murray, 2023). When computing representational alignment between two brain regions, we extracted the off-diagonal elements of the neural matrices and calculated the rank-based correlation between regions. A high correlation means the two regions were representationally similar, which has been suggested to be driven by intensified informational exchange and tighter coupling of time-series between the two regions (see Anzellotti and Coutanche, 2018 for discussion about ’representational connectivity’). As a complementary analysis of representational alignment, we also extracted each region’s fluctuation of BOLD time-series and computed the correlation between brain regions as per the conventional measure of covariation-based functional connectivity.

Following the approach of Ito and Murray (2023), we computed the representational dimensionality for each participant and each brain region. It was calculated as the ‘participation ratio’ that quantified the degree to which all of the dimensions contributed evenly to a region’s neural code. If a region has low dimensionality, the first few dimensions can explain a large proportion of the variance while the remaining dimensions only have negligible contribution. That is, the first few dimensions are enough to capture this region’s overall representation, thus a low participation ratio (only a small proportion of the dimensions have explanatory power). By contrast, if a region has high dimensionality, none of the dimensions are dispensable, and all of them contribute to defining the different aspects of the representation. Using the method of Ito and Murray (2023), the participation ratio was calculated as:

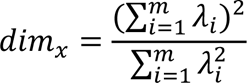

Here *dim_x_* corresponds to the participation ratio of region *X*, and *λ_i_* corresponds to the eigenvalues of the region’s representational distance matrix. As per Ito and Murray (2023), each region’s matrix was decomposed into several eigenvectors/dimensions (the total number of eigenvectors of a matrix is *m*), with eigenvalues (*λ_i_*) quantifying the importance/contribution of a given dimension. The computation was performed individually on each region to quantify whether eigenvalues were disproportionately (low-dimensional) or evenly distributed (high-dimensional) in the whole eigen space (eigenspectrum). Specifically, the more even the eigenspectrum of this region’s matrix, the higher its dimensionality – namely, relying on the initial few eigenvectors are insufficient to explain the full breadth of variance and additional eigenvectors are necessary to explain the full volume of variance, hence driving up the participation ratio. Note, here the feature dimensions were constrained by the number of conditions in our experimental design, rather by the number of voxels in each region, as the goal was to examine how the neural representations differentiated task conditions and to identify the dimensions involved.

We also performed classification analyses on the multivoxel patterns as a complementary approach to RSA. Prior to the classification analyses, we remodelled the data of the RSA experiments in which each block of a task condition was modelled as an individual regressor so as to increase the number of samples for training the classifier in the *k*-fold validation procedure. We performed classification with the Decoding Toolbox (TDT; Hebart et al., 2015), which used a linear support-vector-machine (SVM) for classification with a ‘*C* = 1’ cost parameter (Chang and Lin, 2011). For each classification, a leave-one-run-out 10-fold splitter was used whereby the algorithm learnt to distinguish between relevant categories using the data from nine of the ten runs; its ability to correctly predict was tested using the unseen data from the remaining ‘held-out’ run. This procedure was iterated over all possible combinations of runs used for training and testing. By partitioning the datasets based on different runs of scanning, there was no contamination of information leaking from the training-sets to testing-sets. The accuracy scores were then averaged across folds to produce a mean accuracy score for further statistical analysis. This was done separately for each participant and for each region. Particularly, we used classification to evaluate whether the multivoxel pattern of a brain regions has the ability cross-classify between two disparate domains (e.g., training on the lexical-semantic domain, and then testing on the visual-perceptual or a numerical domain, and *vice versa*).

## Results

We start by providing a roadmap of the analyses conducted: *First*, we began by qualitatively assessing the representational structure for each ROI to identify the resemblances and divergences between the four prefrontal regions, and whether their differences varied in any systematic way along the dorsocaudal-ventrorostral axis. *Second*, correlations/regressions were performed to formally quantify the extent to which the four brain regions’ neural matrices relate to theoretical matrices. In particular, we focused on the comparison among brain regions to explore how the four prefrontal areas differed in their information loading. Comparison among hypotheses was also performed to rank how well the theories characterised the neural patterns. Care was taken to ensure that confounding influences did not contaminate the estimates for each region. *Third*, we investigated ‘representational alignment’ and ‘functional connectivity’ of the four ROIs to understand whether they preferentially communicate with different brain regions beyond the PFC (cf. representational alignment; Ito and Murray, 2023). *Fourth*, we employed machine-learning classification to evaluate how the four ROIs differed in their ability to (*i*) generalise information from one domain to another and (*ii*) differentiate the intricacies of tasks. We then related these machine-learning results with the representational dimensionality of each region to understand whether the state of being high- or low-dimensional systematically varied with the ability to cross-classify or discern details. (cf. Badre et al., 2021 for the links among dimensionality and generalisability).

### Qualitative evaluation on the representational distance matrix

We began by visually inspecting the representational structure of the prefrontal areas’ neural matrices. As evident in Figure 2A, the most obvious information was the gradual emergence of the separation between semantic conditions and non-semantic conditions along the dorsocaudal-ventrorostral axis. Specifically, in the most dorsolateral FEF, there was only mild clustering that differentiated the semantic conditions (i.e., the six conditions in which participants processed semantic meaning) from non-semantic conditions (i.e., the four conditions where participants compared numerical sizes and matched visual shapes). However, the separation strengthened in the MFG, and further manifested in the IFS, and finally culminated in to two distinctly separated clusters in the IFG that contrasted the semantic cluster with the non-semantic one. This bipartite split suggests that, while we manipulated multiple variables of difficulty (easy *vs.* hard), stimuli (words, numbers, shapes), and five task-sets, ‘semantics *vs.* non-semantics’ was the most dominant factor that configured neural representations. This segregation was increasingly clear-cut towards the ventrorostral PFC, hinting a graded change of informational contents couched in the macroscale structure of PFC.

**Figure 2.**
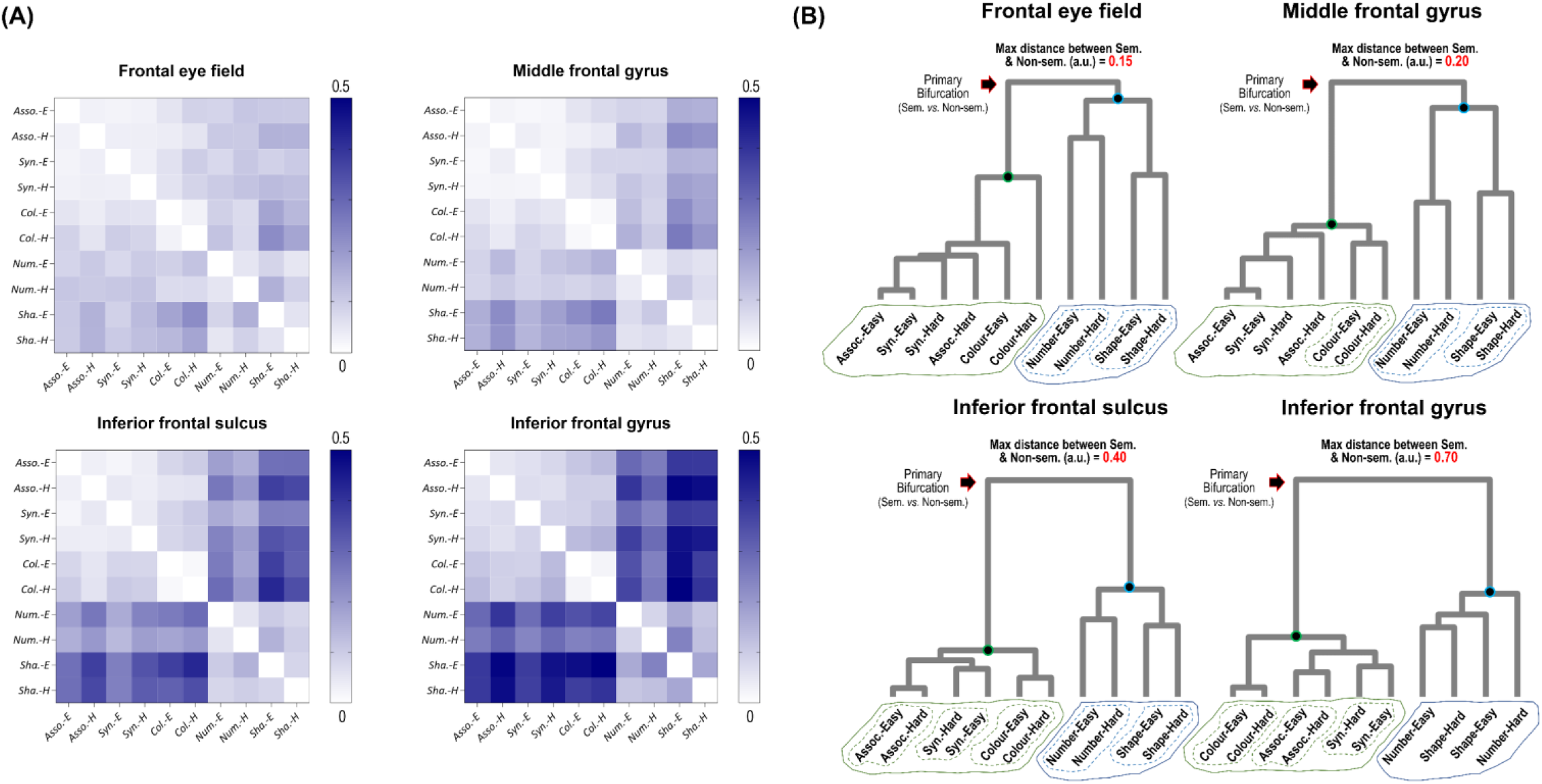
(**A**) The representational distance matrix of the four ROIs. (**B**) Hierarchical clustering was used to analyse the relationship among the neural representations, and dendrograms were used to visualise the relationships. Cophenetic distance (arbitrary unit) was used to quantify the distance between datapoints. It is evident in the dendrograms that the distance between the semantic cluster (the green circle) and the non-semantic cluster (the blue circle) was shortest in the FEF representation, gradually increased in the MFG and IFS, and reached longest in the IFG. Also evident in the figure is that the interposed regions (the MFG and IFS) had more clear-cut representations that individuated the task-sets (the dash-line ovals that enclosed each task) in both semantic and non-semantic domains (the solid-line ovals enclosing each domain), particularly the IFS which individuated all five tasks. By contrast, the FEF and IFG individuated the tasks only in the preferred domain (FEF: non-semantic, IFG: semantic) while conflating the tasks in the non-preferred domain.

We also employed hierarchical clustering to further investigate their representational proximity and visualised the results as dendrograms. As shown in Figure 2B, ‘semantic *vs.* non-semantic’ was the factor driving the primary bifurcation of neural representations, splitting the six semantic conditions from the four non-semantic conditions. While in all the four prefrontal regions this split was seen, closer inspection revealed that the four regions differed in their representational distances between semantic cluster and non-semantic clusters. As Figure 2B shows, this distance (quantified using the Cophenetic coefficient) was shortest in the FEF, gradually lengthened along the MFG and IFS, and was longest in the IFG. This corroborates the qualitative characterisations above, and quantified the degree of segregation. Furthermore, closer scrutiny on the dendrogram branches revealed that the four regions differed in the extent to which the five different tasks were individuated. Specifically, the most dorsocaudal FEF was able to distinguish the two non-semantic tasks (number *vs.* shape) while confusing the three semantic conditions (Association *vs.* Synonym *vs.* Colour knowledge). By contrast, the ventrorostral IFG showed the reverse pattern: it mixed up the two non-semantic tasks, while it was able to clearly discern the three semantic tasks. Crucially, the interposed regions, the MFG and IFS, possessed the ability to tell apart both semantic and non-semantic tasks. Particularly, the IFS encoded each of the five tasks as an individual entity (i.e., the Easy and Hard conditions of the same task were correctly grouped together, while different tasks were independent of one another). Together, these qualitative assessments allowed us to intuitively grasp the overarching pattern that (*i*) the separation of ‘semantic *vs.* non-semantic’ steadily arose along the dorsocaudal-ventrorostral axis and (*ii*) Regions located in the intermediate territory (the MFG and IFS) represented *all* of task-sets as individual entities (i.e., for each task, its Easy and Hard levels were lumped together, due to them sharing the same rules/goals, while segregated from all other tasks). By contrast, regions located in the flanks (the FEF and IFG), could clearly discern tasks of their preferred domain (non-semantic for FEF and semantic for IFG, respectively) while representing tasks in their non-preferred domain in a fuzzy manner.

### Comparing the four prefrontal regions with the theoretically constructed models

We took two approaches to RSA: The first compared the prefrontal subregions and aimed to clarify how they differed in representational content. Thus, the neural matrices were treated as candidates to test against a specific hypothetical matrix. The second treated different hypotheses as candidates to test against the neural matrix of a particular prefrontal region. The two approaches revealed distinct yet complementary information – the first focused on the neural space (the four prefrontal subregions) to compare their goodness of fit with a particular theory while the second focused on the task space (various relationships among task-conditions) to adjudicate which best described the neural pattern.

Prior to conducting RSA, we constructed various hypothetical matrices about the commonality or differences among the task-conditions. As illustrated in Figure 3A, four hypothetical matrices were constructed with each reflecting a unique assumption about the tasks: The *Bipartite* matrix depicted the distinction between semantic and non-semantic domain, stressing the binary segregation between domains while glossing over the differences within each domain. The *Quintipartite* matrix (which means 5-way) only represented the individuality of the five task-sets, stressing the uniqueness of each task while ignoring the difference within a task (i.e., its Easy and Hard levels). The *Complex Structure* matrix represented all graded differences inside the task space – ‘Easy *vs.* Hard’ was nested within each task while the five task-sets were nested within the bipartite split of ‘semantic *vs.* non-semantic’, yielding a graded structure. The *Reaction Time* matrix characterised the pairwise difference of reaction times (RTs) between two conditions – this model was constructed using the behavioural data, taking the absolute value of two conditions’ difference in RTs while ignoring all other aspects of the experimental design; a small difference in RTs meant the two conditions required comparable extent of cognitive effort whereas a large difference meant the two conditions differed greatly in how much effort they taxed.

**Figure 3.**
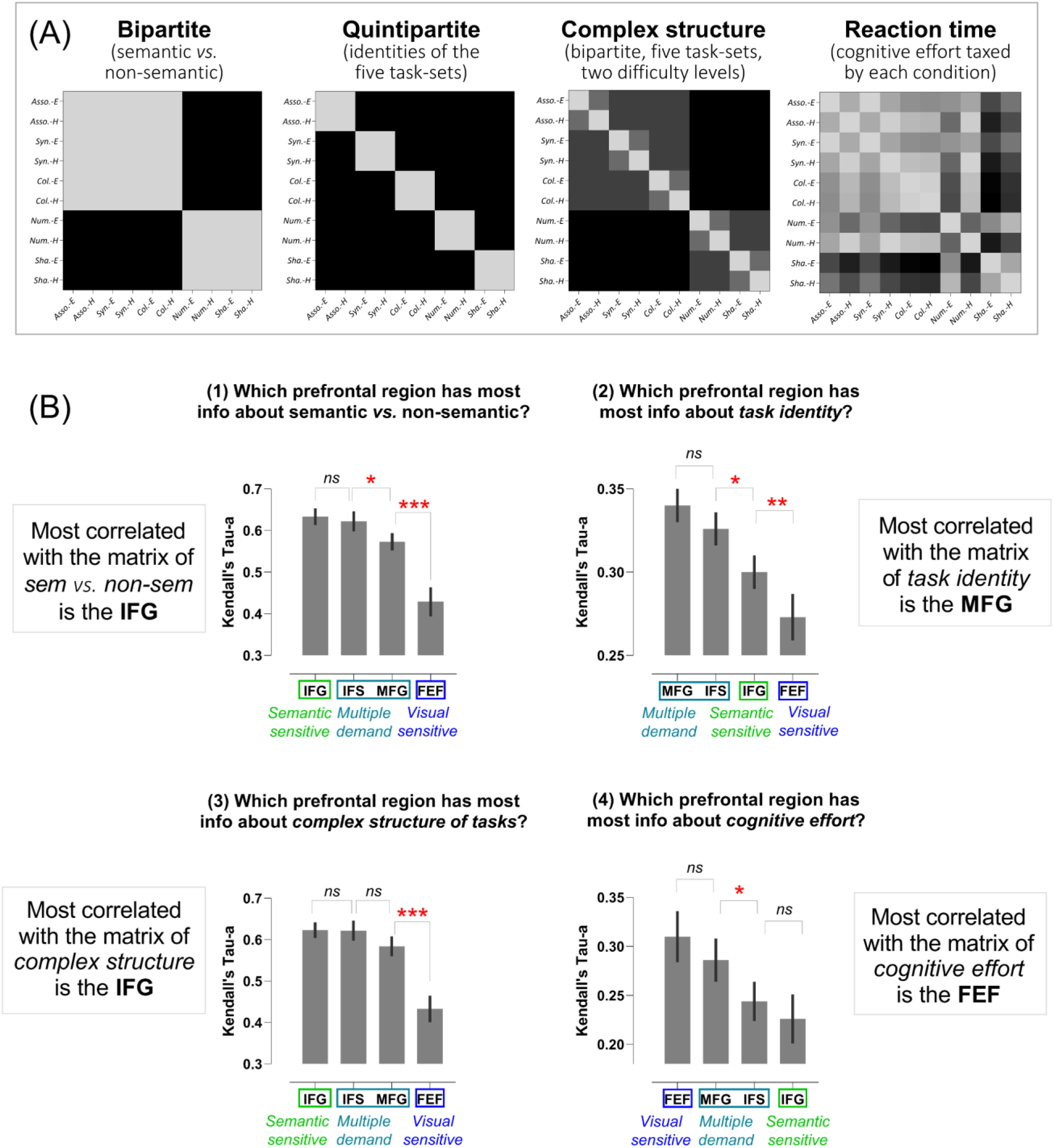
(**A**) The theoretical models set up as the reference matrices of RSA. (**B**) For each model, correlations were computed, associated the four prefrontal regions neural matrices with the reference. Post-hoc comparisons: * *p* < 0.05; *** *p* < 0.001. Error bars are standard error of the mean (SEM).

After setting up the four hypothetical matrices, we began comparing how much the prefrontal regions (as candidates) accorded with each hypothesis. As illustrated in Figure 3B, it is obvious that areas along the dorsocaudal-ventrorostral axis systematically differed in sensitivity to distinct information. Specifically, information about semantic *vs.* non-semantic (namely, the *Bipartite* matrix) was preferentially encoded in the IFG, which steadily decreased dorsally along the axis, and was least encoded in the FEF, yielding robust differences between the regions (*F*_(3,72)_ = 30.39, *p* < 0.001). Information about cognitive effort (the *Reaction Time* matrix) showed the opposite pattern – it was most encoded in the FEF, gradually decreased ventrally along the axis, and least encoded in the IFG (*F*_(3,72)_ = 10.95, *p* = 0.003). Next, information about the individuality of the five task-sets (i.e., the *Quintipartite* matrix) was most encoded in the two intermediate regions (the MFG and the IFS), but dwindled in the dorsal FEF and the ventral IFG (*F*_(3,72)_ = 23.07, *p* < 0.001). Finally, information about the *Complex Structure* of tasks (which considered the nested structure of difficulty levels, task-sets, and semantic *vs.* non-semantic) was encoded by the IFG, IFS, and MFG to comparable extents, but significantly least in the FEF (*F*_(3,72)_ = 33.32, *p* < 0.001). To summarise the results, we found that regions along the dorsocaudal-ventrorostral axis systematically varied in the information ingredients they represented: The dorsocaudal PFC was more sensitive to information about cognitive effort, the intermediate zone was more sensitive to task-sets, and the ventrorostral PFC was more sensitive to the discrepancy of semantic *vs.* non-semantic.

While the results revealed an informational transition in the PFC, control analyses were necessary to preclude the co-variance between regions (collinearity, especially between two contiguous regions) when we estimated the correlative strength unique to each area. Thus, we computed partial correlation (Figure 4) and multiple regression (Figure 5) to assess the unique relationship between each region and each hypothetical matrix; the correlation/regression coefficients were computed individually for each hypothesis, followed by statistical comparisons among the regions. We found partial correlation and multiple regression both replicated the pattern reported above, even after the confounding factors were excluded. Specifically, in partial correlation, we found that the ventrorostral PFC contained more information about semantic *vs.* non-semantic, which gradually decreased dorsally (*F*_(3,72)_ = 12.84, *p* = 0.002); the MFG carried more information about task-sets, which declined in other areas (*F*_(3,72)_ = 4.96, *p* = 0.03); the ventrorostral PFC (particularly the IFG) had more information about the *Complex Structure* matrix, which decreased dorsally (*F*_(3,72)_ = 10.19, *p* = 0.004); the dorsocaudal PFC (particularly the FEF) possessed more information about behavioural effort, which reduced ventrally (*F*_(3,72)_ = 13.78, *p* = 0.001). A highly consistent pattern was observed in the data of regression analysis: the IFG’s neural pattern best fitted the matrix of ‘semantic vs. non-semantic’, with the *β* coefficients gradually decreasing in dorsal regions (*F*_(3,72)_ = 9.60, *p* = 0.005); the IFG pattern also best fitted the matrix of *Complex Structure*, with the *β* coefficients steadily reducing in dorsal areas (*F*_(3,72)_ = 8.72, *p* = 0.007); the dorsocaudal PFC (particularly the FEF) best fitted the matrix of *Reaction Time*, with the *β* values gradually waning in ventral regions (*F*_(3,72)_ = 14.29, *p* = 0.001); finally, we only observed a non-significant weak trend in the regression model for the hypothesis of task identity (*F*_(3,72)_ = 3.04, *p* = 0.09) – the middle PFC (the MFG and IFS) had numerically higher *β* values than the dorsal FEF and ventral IFG. Taken together, the results of partial correlation and multiple regression replicated the overall pattern found in the unadjusted correlation – there is graded difference between regions, with behavioural effort preferentially encoded in the dorsocaudal PFC, task-sets in the middle PFC, and abstract structures (e.g., semantic *vs.* non-semantic) in the ventrorostral PFC.

**Figure 4.**
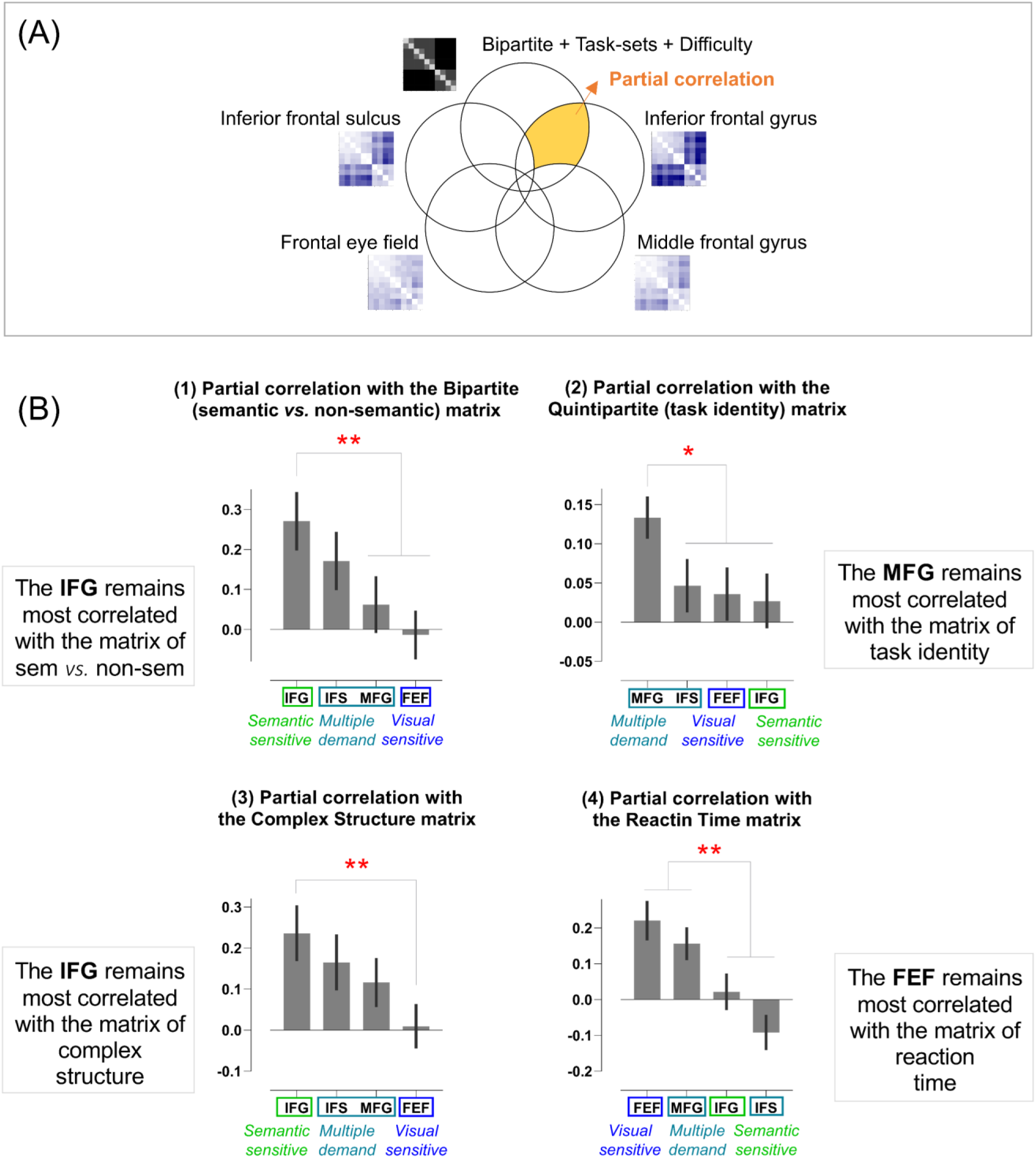
(**A**) This example illustrates partial correlation in a Venn diagram. The unique relationship between the IFG and the *Complex Structure* hypothesis (which is what partial correlation identifies) is shaded in yellow while the co-variance driven by other areas are excluded. (**B**) For each hypothesis, we computed its partial correlations with each of the four prefrontal areas, followed by statistical tests comparing the four regions. Post-hoc comparisons: ** *p* < 0.005; * *p* < 0.05. Error bars indicate SEM.

**Figure 5.**
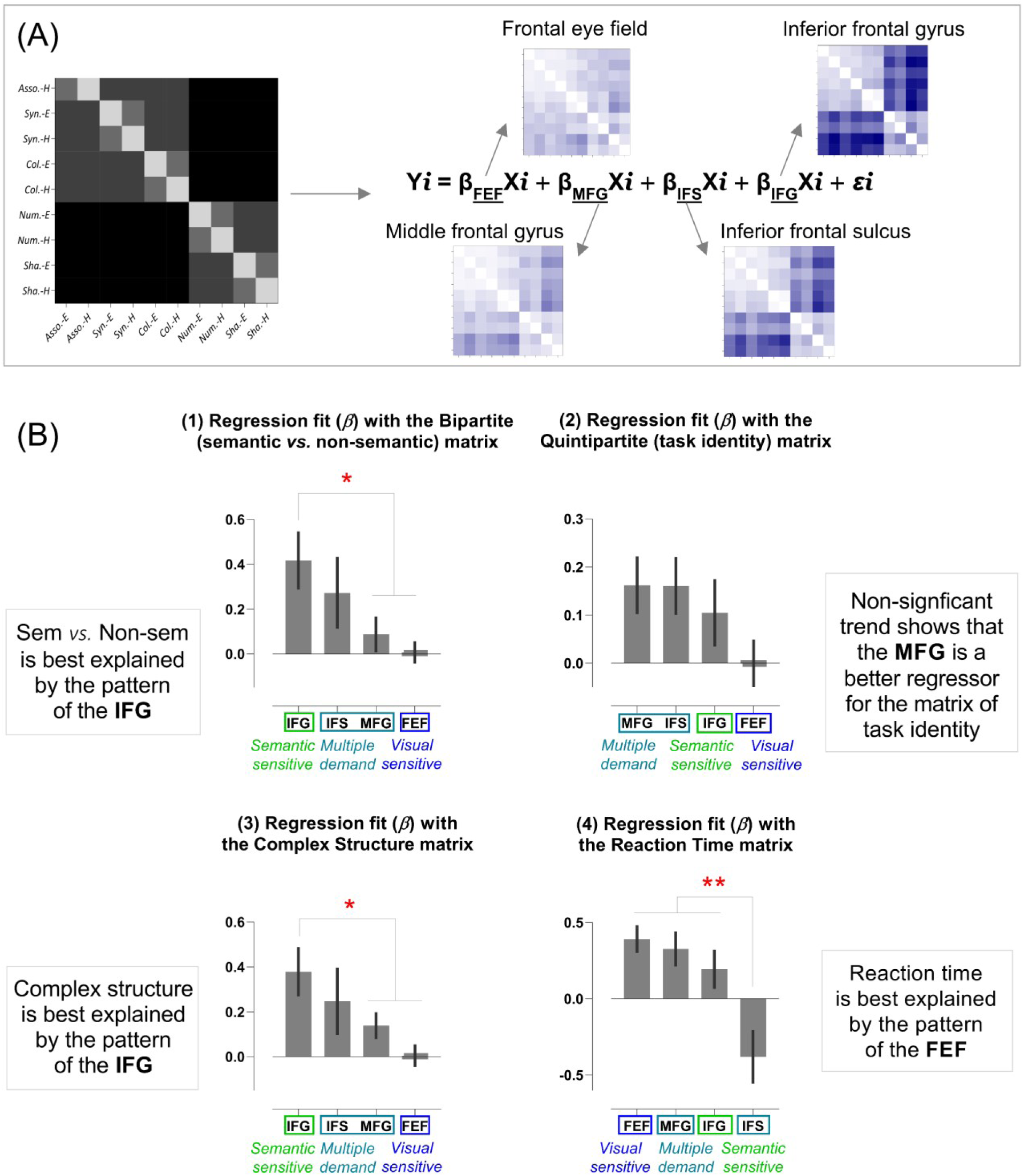
(**A**) In this example of regression analysis, the four prefrontal regions’ neural patterns are included as regressors to predict the matrix of Complex Structure hypothesis. Regression fit (*β* weight) indicates the unique explanatory power of a region when the effects of other regions are held constant. (**B**) For each hypothesis, the *β* weight associated with each regressor were computed, followed by statistical comparisons among the regions. Post-hoc comparisons: ** *p* < 0.005; * *p* < 0.05. Error bars indicate SEM.

In addition to comparing the prefrontal regions to know which best fitted with a certain hypothesis, we also carried out the inverse test – comparing different hypotheses to know which best described a region’s neural pattern. For this analysis, we additionally constructed the *Tripartite* matrix (3-way division in the task space) and *Quadripartite* matrix (4-way division) but excluded the *Complex Structure* matrix (owing to its multicollinearity with other matrices), yielding five hypotheses as candidates (Figure 6A). Partial correlation was calculated to quantify the unique relationship between a region and a target hypothesis while excluding the influences from all other hypotheses. As shown in Figure 6B, for each of the four regions, the five hypotheses differed significantly in the strength of association with the neural data (FEF: *F*_(4,96)_ = 41.49, *p* < 0.001; MFG: *F*_(4,96)_ = 53.91, *p* < 0.001; IFS: *F*_(4,96)_ = 61.18, *p* < 0.001; IFG: *F*_(4,96)_ = 125.07, *p* < 0.001). We found that, the *Bipartite* matrix (semantic *vs.* non-semantic) was generally the best candidate to account for the neural data, significantly outperforming all other hypotheses (with the exception that, in the FEF, *Reaction Time* was on a par with the *Bipartite* matrix). These statistically support our interpretation of Figure 2A that ‘semantic *vs.* non-semantic’ predominantly determines how clustering was formed.

**Figure 6.**
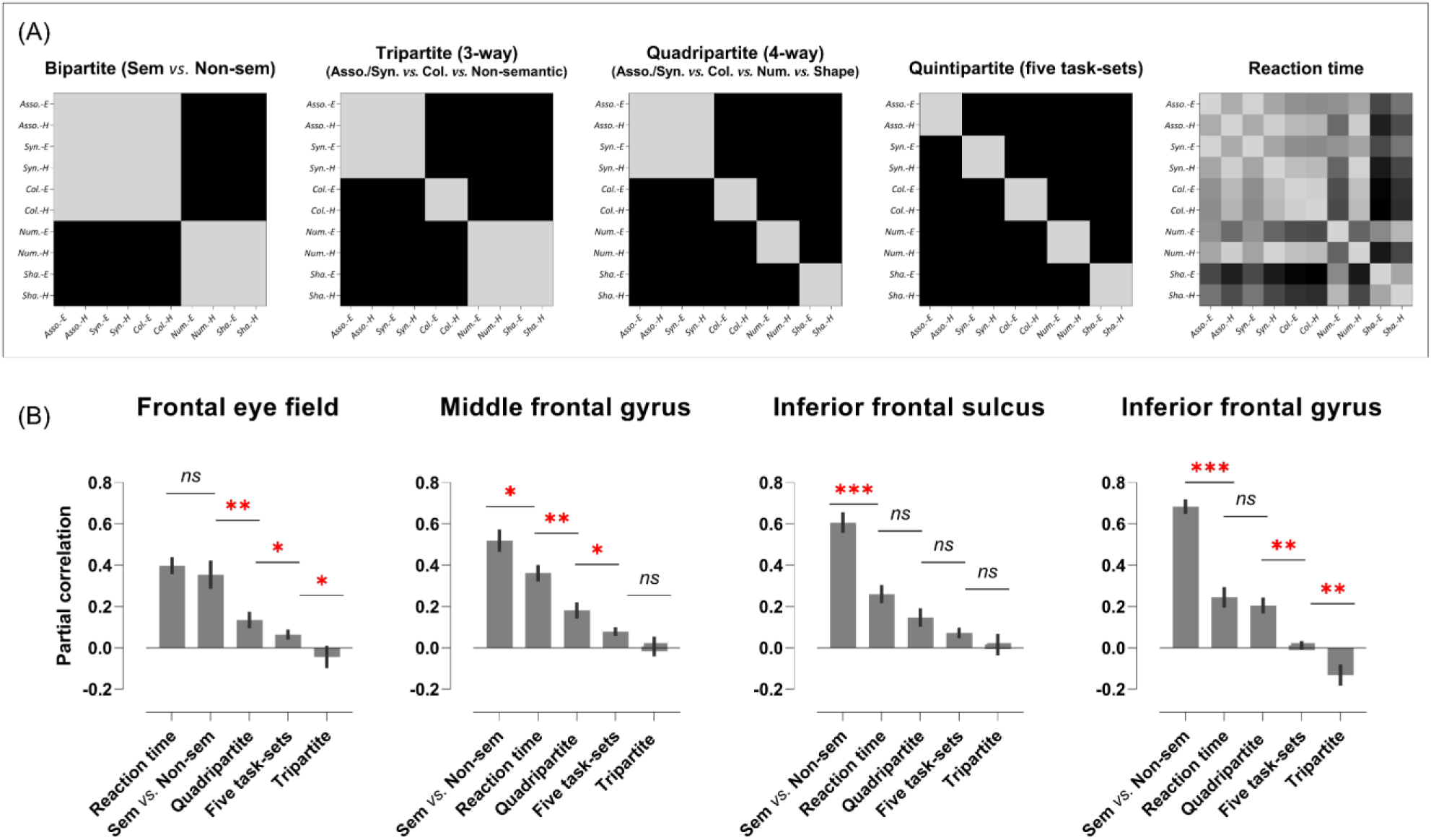
(**A**) Five theoretical matrices were constructed as the candidates to assess their relationship with each region’s neural pattern. (**B**) The partial correlation between the theoretical matrices and each of the four regions. Post-hoc comparisons: *** p < 0.001; ** *p* < 0.005; * *p* < 0.05. Error bars indicate SEM.

Taken together, the results revealed that, for all of the prefrontal regions, ‘semantic vs. non-semantic’ was the predominant factor that configured neural representations relative to every other hypothesis. However, when specifically comparing the four regions to investigate how they fitted each hypothesis, we found robust differences among regions such that distinct types of information was preferentially registered in different subparts along the dorsocaudal-ventrorostral axis of the PFC.

### Representational alignment with regions beyond the prefrontal cortex

It has been established in studies of both monkey cytoarchitecture (e.g., Saleem et al., 2014) and human neuroimaging (e.g., Jung et al., 2022) that different subdivisions in the PFC tend to connect with separate regions outside the PFC. The dorsal PFC, especially the FEF, is preferentially connected with the posterior parietal cortex (PCC) and jointly form a circuit heavily involved in spatial attention (Yeo et al., 2011; Xu, 2018), whereas the ventral PFC, especially the IFG, is preferentially connected with the posterior middle temporal gyrus (pMTG) and they jointly form a circuit heavily involved in retrieving semantic memory (Davey et al., 2016; Chiou et al., 2018; Jackson, 2021). We speculated that connectomic fingerprints (i.e., preferential linkage to different areas, cf. Mars et al., 2018) might be the mechanism that drives distinct information to be preferentially registered in different parts of the PFC (in line with the broader hypothesis that local processing is shaped in part by the pattern of its connectivity with the rest of the brain that constrains this area’s input/output). To investigate this, we employed representational alignment analysis (cf. Ito and Murray, 2023) and compared the results of representational alignment with seed-based functional connectivity. A high correlation between two regions’ representational matrices indicates that information encoded by the two regions are ‘configurationally aligned’ and is interpreted as more informational exchange between the two areas (greater connectivity strength; cf. Anzellotti and Coutanche, 2018).

As shown in Figure 7A, the neural pattern of pMTG was most aligned with that of the IFG (i.e., highest resemblance), followed by the IFS and MFG, and was least aligned with the FEF (*F*_(3,72)_ = 74.62, *p* < 0.001). This pattern was mirrored in the result of functional connectivity – the timeseries of pMTG, as the seed, was most synchronised with that of the IFG, followed by the IFS and MFG, and least with the FEF (*F*_(3,72)_ = 51.32, *p* < 0.001). This gradient (which peaked in the ventral PFC and gradually weakened dorsally) was flipped when we examined the four regions’ relationship with the PCC (see Figure 7B): The PCC’s neural pattern was representationally most similar to the FEF, followed by the MFG and IFS, and least similar to the IFG (*F*_(3,72)_ = 8.62, *p* = 0.007); also, the PCC’s timeseries was most in synchrony with the FEF, followed by MFG and IFS, and least with the IFG (*F*_(3,72)_ = 87.32, *p* < 0.001). Together, these results replicated the functional and structural divide of the dorsal *vs.* ventral PFC, echoing the seminal classic study of Goldman-Rakic (1984), and revealed a parallel between representational similarity and functional connectivity.

**Figure 7.**
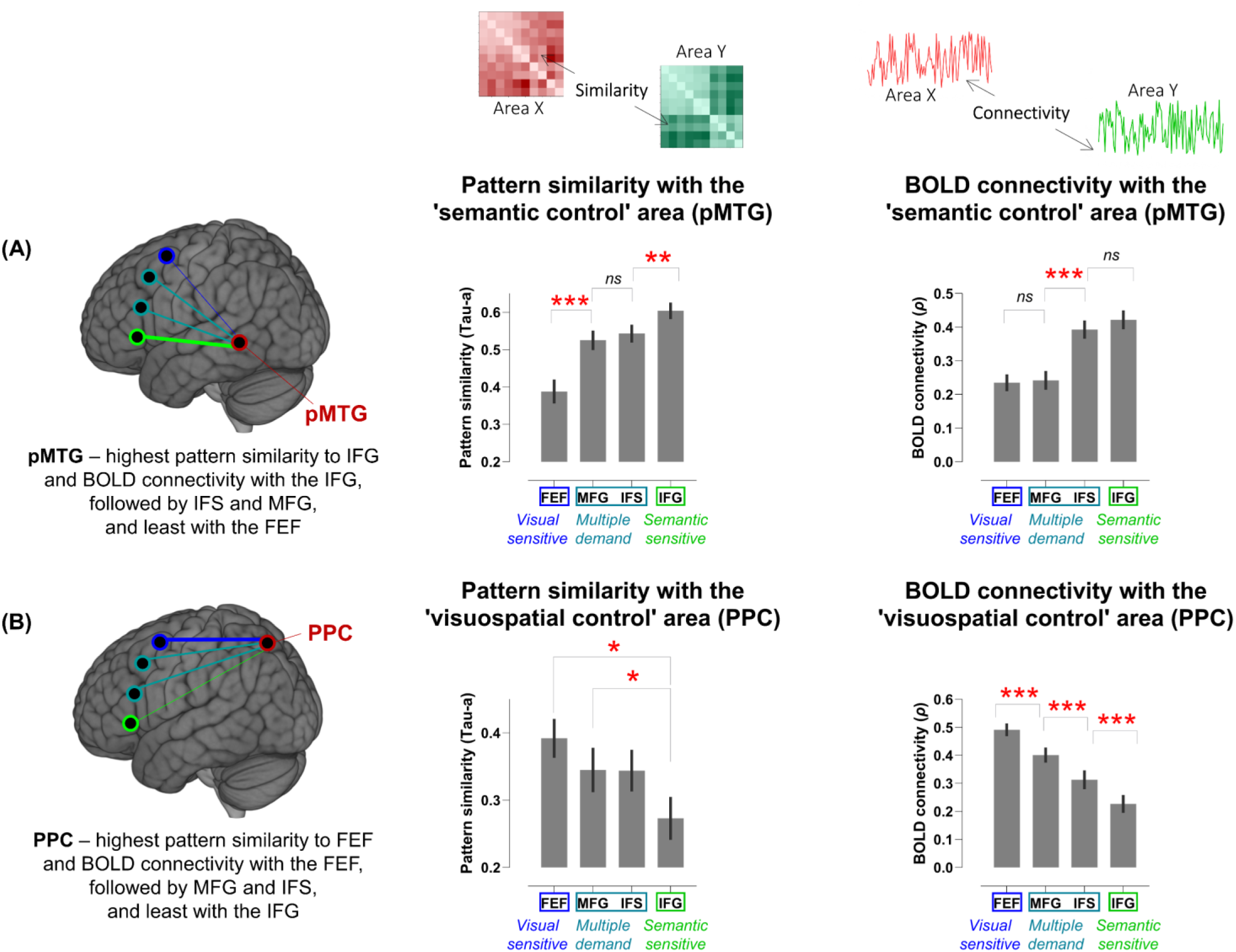
(**A**) Representational alignment: Comparing the similarity of neural patterns and the strength of functional connectivity between the pMTG (a region heavily involved in semantic control) and the four prefrontal regions. (**B**) Functional connectivity: Comparing the similarity of neural patterns and the strength of functional connectivity between the PPC (a region heavily involved in visuospatial attention) and each of the four prefrontal regions. Post-hoc comparisons: *** *p* < 0.001; ** *p* < 0.005; * *p* < 0.05. Error bars indicate SEM.

### Dimensionality, generalisability, and expressivity of neural representations

Dimensionality-reduction techniques, when applied to multivariate neural data, enable researchers to dissect informational content and extract major dimensions (Rigotti et al., 2013; Brincat et al., 2018; Bartolo et al., 2020). A key approach is to quantify the dimensionality of neural representations (Badre et al., 2021; Ito and Murray, 2023). In technical terms, the dimensionality of a brain region’s representation refers to the number of variables needed to capture the essence of its representation. Research has shown that high dimensionality of neural codes is associated with high expressivity (i.e., the region can represent subtle differences, and is therefore highly expressive) but it comes with the cost of low generalisability (the neural code captures too much peculiarity about each stimulus, hurting its ability to generalise across contexts – akin to overfitting). By contrast, low dimensionality is associated with low expressivity (unable to tell apart subtle differences) but high generalisability (i.e., the neural representation captures the broad-stroke information, which can be extracted from one context and applied to another) (see Badre et al., 2021 for the trade-off among dimensionality, expressivity, and generalisability).

We investigated how representational dimensionality varied between the four prefrontal regions, and related dimensionality with generalisability and expressivity. To begin with, we examined whether neural representations of the four regions differed in their generalisability across contexts. This was achieved with a multivariate cross-classification: the algorithm was trained to classify Easy *vs.* Hard on the data of a semantic task and tested using the unseen data of a non-semantic task (and *vice versa*). Successful decoding means the neural code of a region contained generalisable information about difficulty across domains. As shown in Figure 8, we found that the generalisability systematically varied along the dorsocaudal-ventrorostral axis such that the accuracy of cross-domain decoding of difficulty was highest in the FEF, declined in the MFG and IFS, and became non-existent in the IFG. This was found in the cross-classification between the Association task and non-semantic task (*F*_(3,72)_ = 15.66, *p* < 0.001), and between the Synonym task and non-semantic (*F*_(3,72)_ = 12.98, *p* < 0.001), unaffected by whether the non-semantic task was Number or Shape (both *F*s < 1). A similar albeit non-significant trend (highest in the FEF and gradually dropping in ventral regions) was seen in the results of the Colour Knowledge task (*F*_(3,72)_ = 2.91, *p* = 0.11). Next, we computed the dimensionality for each of the four prefrontal regions, using the methods of Ito and Murray (2023; see Methods for the mathematical details). As shown in Figure 9A, representational dimensionality was highest in the IFG, followed by the IFS and MFG, and dropped to lowest in the FEF (*F*_(3,72)_ = 24.64, *p* < 0.001). Crucially, there is an inverse relationship between dimensionality and generalisability – a trade-off between these two features. As shown in Figure 9B (the Association task) and 9C (the Synonym task), the IFG showed highest dimensionality yet lowest generalisability, which could be understood as the neural patterns of IFG being engrossed by the high-dimensional task features and failing to identify any generalisable patterns across contexts. By contrast, the FEF exhibited lowest dimensionality but highest generalisability, suggesting that this region used ‘stereotyped’ patterns to represent Easy *vs.* Hard (which was shared across the five tasks included, thus enabling cross-domain classification).

**Figure 8.**
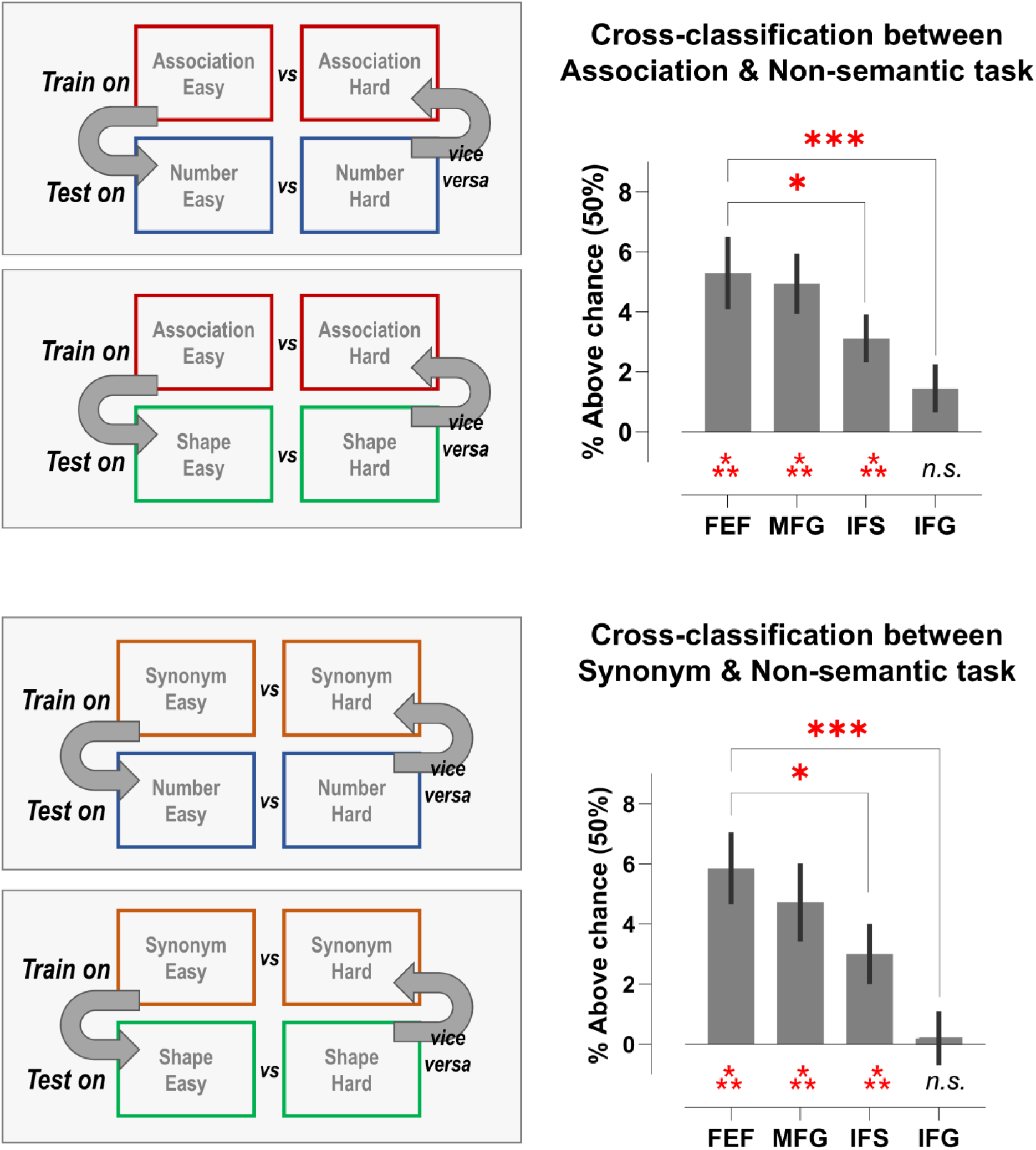
In the multivoxel cross-classification, the support-vector machines were trained to classify Easy *vs.* Hard of a semantic task (either the data of Association task or that of the Synonym task) and were tested with a non-semantic task (either the Number task or the Shape task), and *vice versa* – trained on non-semantic, tested on semantic. Chance-level was 50% in this binary classification. Because the outcome was unaffected by whether the cross-classification was against the Number task or the Shape task (*F*s < 1), the averaged results were shown here to highlight the main effect of ROIs that cross-classification accuracy was high in the dorsal PFC and dropped steadily in the ventral PFC. Post-hoc comparisons: *** *p* < 0.001; ** *p* < 0.005. Error bars indicate SEM.

**Figure 9.**
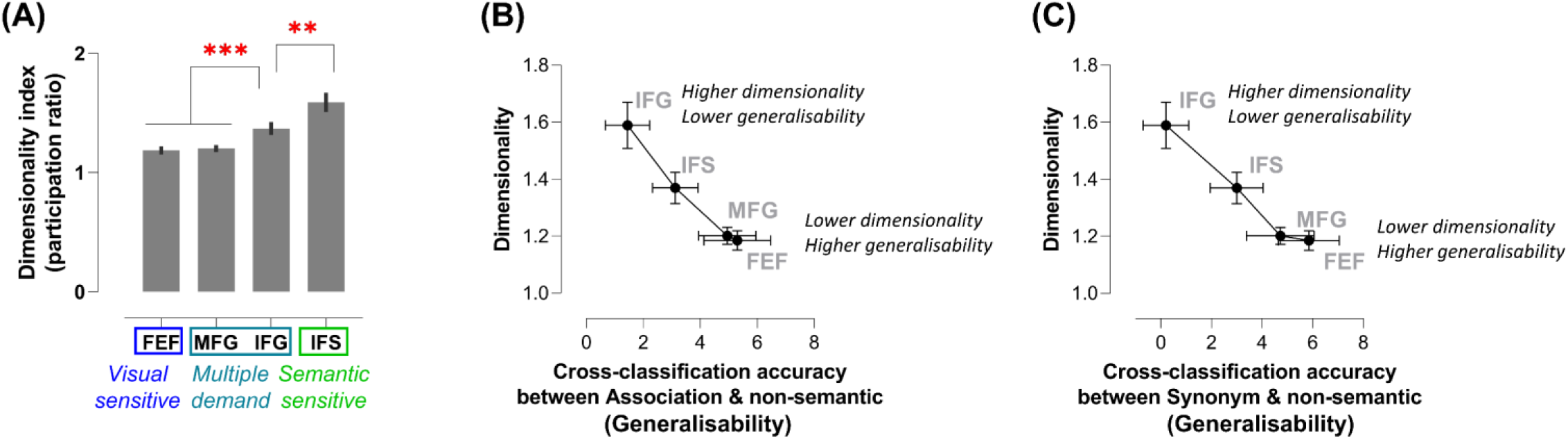
(**A**) Dimensionality index, computed as the participation ratio of the eigenvariables of a region’s representational matrix, of the four prefrontal regions. We found that dimensionality has an inverse relationship with cross-domain classification accuracy (which was a proxy measure of generalisability, indicating how well the neural pattern could generalise information from one domain to another). This was found in the results of the Association task (**B**) and Synonym task (**C**) such that highly dimensional regions tended to exhibit lower generalisability. Post-hoc comparisons: *** *p* < 0.001; ** *p* < 0.005. Error bars indicate SEM.

We also analysed the relationship between dimensionality and expressivity. According to Badre *et al*. (2021), highly expressive neural representations encode minute differences between task-conditions, easily separating them from each other, while low expressive representations fail to tell the difference. Based on this definition, we performed a 10-way classification for each of the four prefrontal regions to measure expressivity, testing the ability to distinguish details across the entire scope and predict the one correct label from 10 task-conditions. Thus, decoding accuracy served as a proxy measure of expressivity to assess how much nuance was discernible by the representation. As Figure 10A shows, decoding accuracy was highest in the IFG, gradually declined dorsally, and became lowest in the FEF (*F*_(3,72)_ = 56.69, *p* < 0.001). As shown in Figure 10B, decoding accuracy had a positive relationship with dimensionality such that the most expressive region (the IFG) had highest dimensionality while the least expressive region (the FEF) had lowest dimensionality.

**Figure 10.**
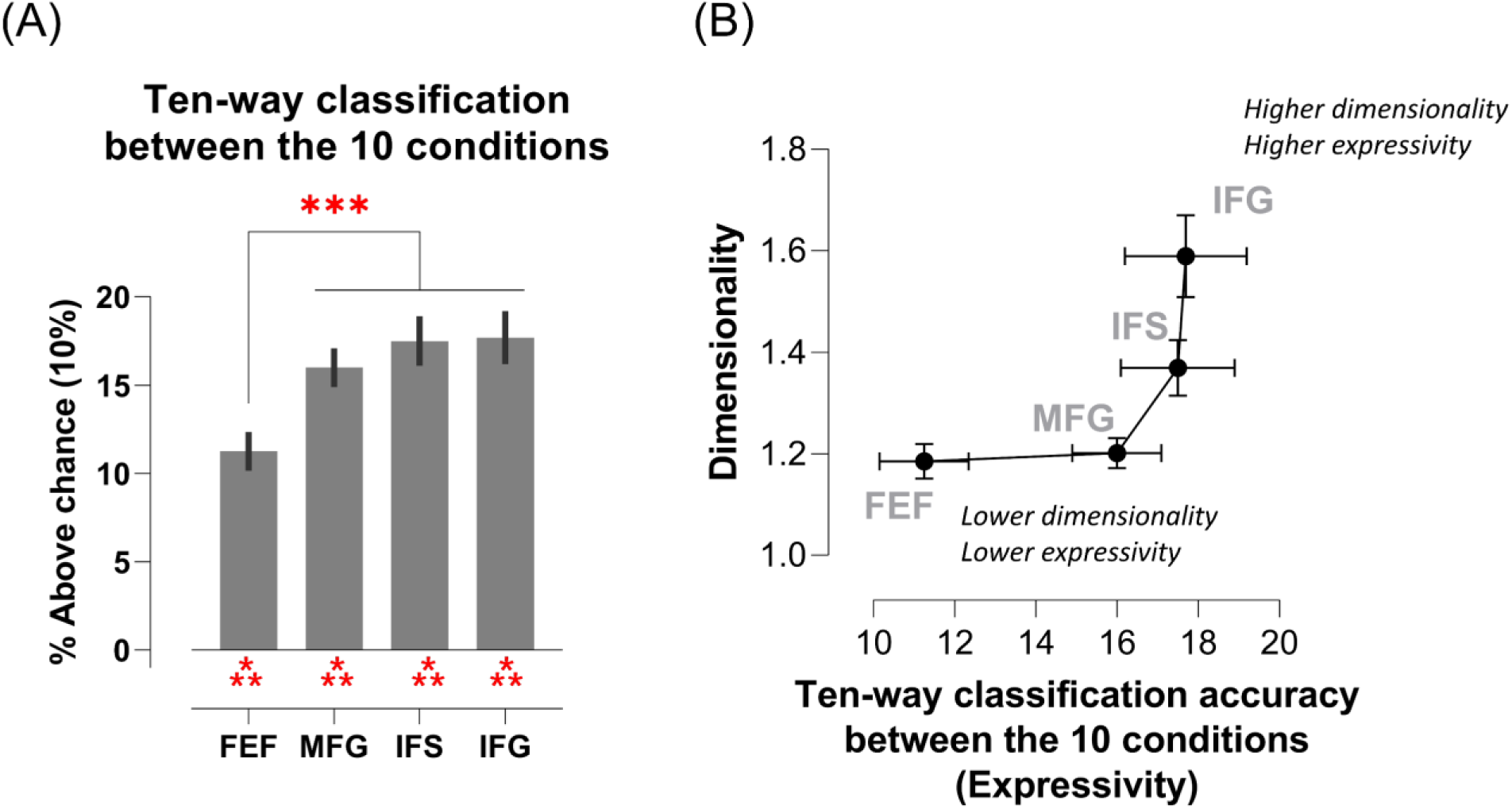
(**A**) In the 10-way cross-validated classification, the support-vector machines were trained to predict the correct condition label from 10 possible choices. This 10-way classification required more details about the differences between conditions than, for example, the binary classification of semantic *vs.* non-semantic, and served as a proxy quantifying a region’s representational expressivity. Chance-level was 10% in this 10-way classification. The accuracy of such fine-grained classification was higher in the ventral PFC and lower in the dorsal FPC. (**B**) There was a positive relationship between dimensionality and expressivity that highly dimensional regions (e.g., the IFG) tended to show higher expressivity. Post-hoc comparisons: *** *p* < 0.001. Error bars indicate SEM.

Taken together, our investigation into the relationship among dimensionality, generalisability, and expressivity revealed a trade-off between these features, concurring with the proposal of Badre *et al*. (2021). Moreover, all these features varied gradually along the dorsocaudal-ventrorostral axis of PFC, with representations being, respectively, most generic (low-dimensional) in the dorsocaudal PFC and most specific (high-dimensional) in the ventrorostral PFC.

## Discussion

In the present study, we designed a multifarious paradigm in which participants performed three types of semantic tasks and two types of non-semantic tasks, with difficulty level manipulated for each task. This design merged multiple variables into a multifaceted structure and enabled us to leverage RSA to identify the abstract dimensions that defined neural representations during task performance and verify whether distinct information was preferentially encoded in different subparts of the PFC. Results showed a graded transition along the dorsocaudal-ventrorostral axis: The FEF was sensitive to the information about cognitive effort, preferentially connected with the dorsal parietal lobule, encoding task difficulty in a representationally generalisable fashion yet dimensionally impoverished. By contrast, the IFG was sensitive to the complex/abstract structure of our task paradigm (particularly to the principal division of semantic *vs.* non-semantic), preferentially connected with the left MTG, representationally expressive to distinguish subtleties, and dimensionally richest among the four areas. Finally, the areas interposed in the middle zone (the MFG and IFS) were sensitive to the individuality of task-sets, ranked in the middle in terms of their connectivity, dimensionality, generalisability, and expressivity. Together, by leveraging subject-specific localisation to capture the idiosyncrasy of individual activation (Fedorenko, 2021) and RSA to infer informational contents (Freund et al., 2021a), we showed how highly dimensional prefrontal representations could be decomposed and compared with various hypothetical constructs about cognitive control, revealing the gradual transformation of task-relevant information in different PFC subparts.

Below we expound on four issues: (1) How do the present findings relate to previous literature about semantic *vs.* non-semantic processing in the PFC; (2) How can dimensionality explain the functional repertoires of PFC subregions; (3) Functional connectivity and the emergent prefrontal gradient; (4) Different approaches to study the organisation of cognitive control in the PFC.

### Semantic control system vs. multiple-demand system in the PFC

Primarily based on univariate analysis, a dorsal-ventral divide has been established in the literature: The ventrolateral PFC is robustly involved in various goal-directed operations on semantic meaning (e.g., Thompson-Schill et al., 1997; Badre et al., 2005; Chiou et al., 2018; Gao et al., 2021; Zhang et al., 2021; Chiou et al., 2022) and joins forces with the pMTG as the ‘semantic control’ system (e.g., Davey et al., 2016; Alam et al., 2019; Jackson, 2021). Rather than specific to semantic control, the ventrolateral PFC is also engaged by other inwardly-directed cognitive tasks (e.g., episodic memory: Vatansever *et al*., 2021; autobiographical memory: Chiou *et al*., 2020; language comprehension: Fedorenko *et al*., 2011), and it tends to be suppressed by outwardly-directed tasks (e.g., visual search: Chiou *et al*., 2020). However, instead of simply preferring inward cognition, the ventrolateral PFC is also sensitive to executive demand, as its activation increases with the difficulty of semantic processes (but not the difficulty of outward cognition; Humphreys and Lambon Ralph, 2017; Chiou et al., 2022). This selective and combinatorial profile (e.g., preferring semantic tasks especially when it is difficult) reflects a fusion between semantics and executive control. In contrast, the ‘multiple-demand’ system is sensitive to executive demand on outwardly-oriented perceptual/motoric tasks (Duncan, 2010) and occupies a set of widely distributed cerebral patches in the prefrontal and parietal lobes (Assem et al., 2020; Assem et al., 2021). In the lateral PFC, the ‘semantic control’ system is located in subregions ventral to the ‘multiple-demand’ system. Advancing from univariate data, our multivariate decoding lends support to this dorsal-ventral divide and further reveal the informational contents. For example, transitioning from dorsocaudal to ventrorostral PFC, the preferred contents vary from cognitive effort, via task-sets, to semantics meaning, consistent with a progression from ‘multiple-demand’ regions to ‘semantic control’ regions. Furthermore, our dimensionality analysis showed that IFG representations had higher dimensionality yet lower generalisability than dorsal regions. The high dimensionality in ventral regions is consistent with a non-linear profile of responses (the IFG cannot be defined simply by either difficulty or semantic as it prefers *both* easy mind-wandering and difficult semantic tasks), whereas the low dimensionality in dorsal regions fits with their straightforward profile of responses (i.e., ‘multiple-demand’ regions prefer difficulty processes, irrespective of tasks, stimuli, or modality).

### Dimensionality in different prefrontal regions

It has been argued that, to precisely encode the world and efficiently respond to novel challenges, the brain needs to balance precision (truthfully representing details) against efficiency (swiftly reusing learnt skills to tackle similar problems) Hence, high- and low-dimensional representations are both necessary to enable such adaptive behaviour (Fusi et al., 2016; MacDowell et al., 2022). To represent complex thoughts and actions, high-dimensional representations are needed to capture all minutiae; to efficiently adapt to new situations, low-dimension representations are needed to encode a ‘sketch’ of the novel challenge, identify its similarity with former situations, and generalise previous solutions. As suggested by Badre *et al*. (2021), there is a computational trade-off as the brain balances between low and high dimensionality. This fits with our findings: The ventrorostral PFC was high-dimensional, which enabled distinguishing subtleties yet compromised its generalisability across domains, whereas the dorsocaudal PFC was low-dimensional, which allowed extrapolating from one domain to predict another domain but its coding was less elaborate. Critically, dimensionality steadily varied along the prefrontal gradient, which may be driven by graded changes of connectivity with posterior regions (which we discuss next).

### Connectivity and functional heterogeneity in the PFC

It has been established that the dorsal parietal and ventral temporal pathways of the visual system are preferentially connected with, respectively, the dorsolateral and ventrolateral PFC. This was initially identified by the work of Goldman-Rakic using monkey electrophysiology (Goldman-Rakic, 1984) and was later replicated in human neuroimaging (e.g., Shmuelof and Zohary, 2005), as well as in the results of representational alignment and seed-based connectivity that we report in the present study. Recent evidence further show that the difference extends beyond the visual system (Haber et al., 2022; Jung et al., 2022), with the dorsolateral PFC more connected with regions for attentional control and the ventrolateral PFC more connected with regions for semantic memory. As argued by Mars *et al*. (2018), each region’s unique pattern of input and output connections shapes its functional repertoire, driving functional heterogeneity consequently. In fact, not only the PFC, connectivity has been shown to cause functional diversity within the temporal cortex (Jackson et al., 2016; Bajada et al., 2017), within the parietal cortex (Humphreys et al., 2022), and within the occipital cortex (Konkle and Caramazza, 2017). Taken together, robust evidence suggests that connectivity is a major factor that constrains how functions arise from brain structures.

### Different approaches to study the organisation of cognitive control in the PFC

Different approaches used to study the functional organisation of cognitive control in the PFC often yield different conclusions. One approach is parcellating the PFC into a patchwork of cortical parcels (Glasser et al., 2016) and identifying the parcels preferring harder to easier tasks (Assem et al., 2020; Assem et al., 2021). This approach reveals five patches in the PFC, which topographically form a circular arrangement that constitutes the core areas for cognitive control in the PFC. Another approach assumes that cognitive control is hierarchically organised along the dorsocaudal-ventrorostral axis of PFC (for review, see Badre and Nee, 2018; Badre and Desrochers, 2019). This approach typically hierarchically manipulates the contingency between stimuli and responses that higher-order contexts stipulate different responses to the same stimuli (namely, as the superordinate/abstract contexts vary, the concrete correspondences between stimuli and responses also vary; see Badre and D’Esposito, 2007; Nee and D’Esposito, 2016; Nee, 2021). With such hierarchically nested structures in the design, graded changes in the PFC have been found – the dorsocaudal division is involved in executing action, the middle division is involved in maintaining task-sets, and the ventrorostral division is involved in representing a superordinate schema of task contexts (e.g., Nee and D’Esposito, 2016). It is, however, important to note that arbitrating on the various approaches is difficult given the drastic differences in experimental designs, ways to parcellate the PFC, and analytical methods. Functional dissociations in the PFC can vary depending on a wide range of factors – the choices of tasks being compared (e.g., hierarchical or ‘hard *vs.* easy’ contrast), how ROIs are defined (coordinate-based, parcel-based, or using a localiser to select voxels within a scope), and the analysis performed (univariate, multivariate, connectivity, etc.). Thus, it is unsurprising that they have yielded disparate conclusions.

### Limitation of the present study and future directions

Given the differences in task design, loci of the ROIs, and analytical methods, it is difficult to establish the correspondence between the PFC gradient we identified and those reported previously. Ostensibly, we found a gradient from behavioural effort, via task-sets, to semantics, which is morphologically consistent with the ‘concrete-to-abstract’ transition reported previously (e.g., Badre and Nee, 2018). However, future research is needed to closely match experimental parameters, potentially adopting a hierarchical-nested design that incorporates both semantic and non-semantic elements and using an individual localiser to select voxels within parcels of the high-precision fine-grained parcellation of the Human Connectome Project. With all these parameters incorporated, it will hold promise for reconciling discrepant findings and clarifying how cognitive control is implemented in the PFC.

## Acknowledgements

This research was funded by an MRC programme grant and intramural funding to MALR (MR/R023883/1; MC_UU_00005/18), and a Sir Henry Wellcome Fellowship (201381/Z/16/Z) to RC.

## Notes

**Conflict of interest:** The authors declare no competing financial interests.

### Competing Interest Statement

The authors have declared no competing interest.

