## Extended data (Figure 1-1) for "The dimensionality of neural coding for cognitive control is gradually transformed within the lateral prefrontal cortex"

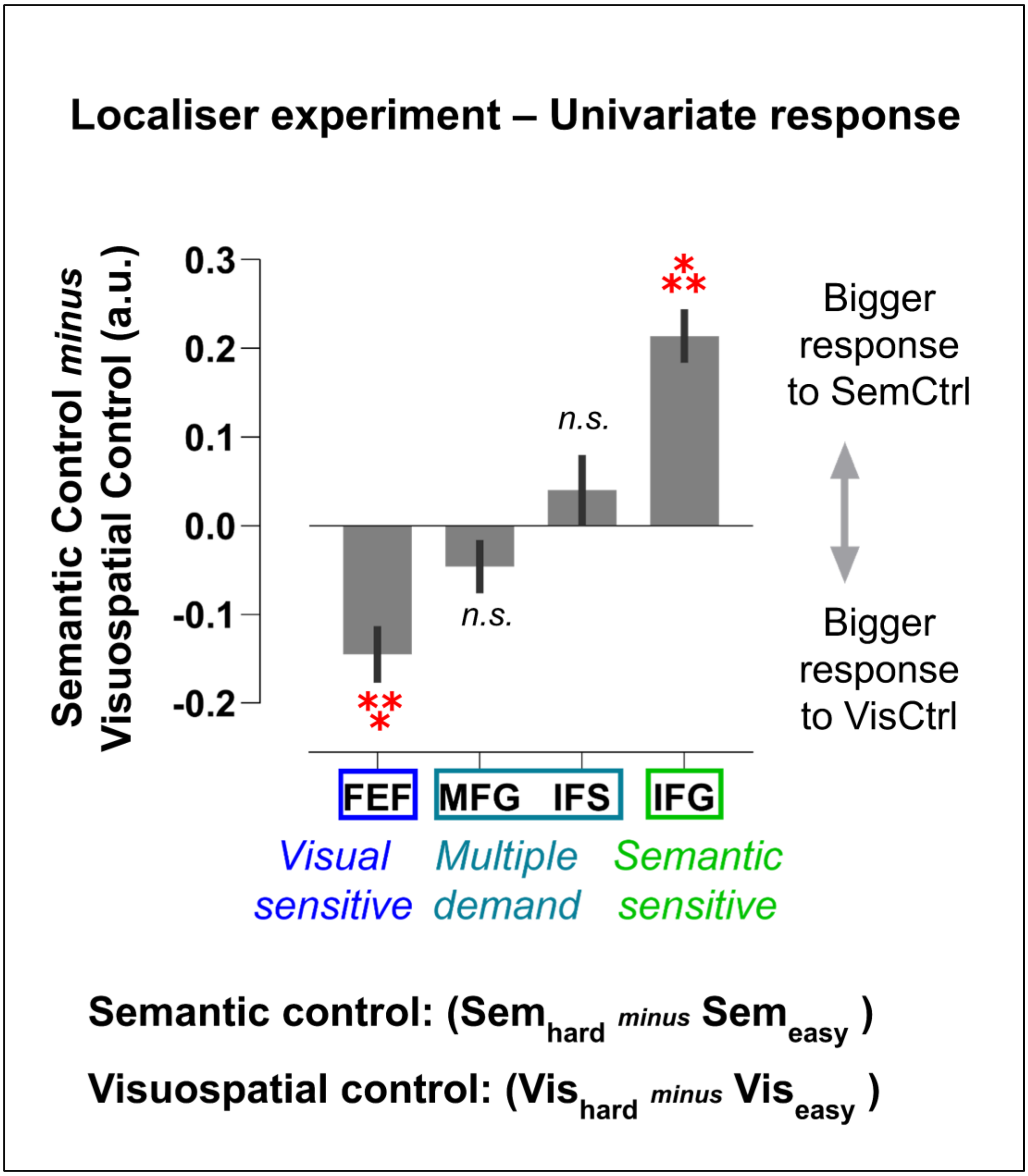


**Figure 1–1**. Note, the data used in this supplementary analysis are based on the localiser experiment, rather than the RSA decoding experiment. In this analysis, we examined the univariate responses of the prefrontal ROIs to semantic control (Semantic-Hard > Semantic-Easy) *vs.* visuospatial control (Visuospatial-Hard > Visuospatial-Easy), and then we compared between the four ROIs to understand whether they exhibited any preference for semantic control or visuospatial control.

As illustrated in Figure 1–1, we found a significant main effect of ROIs ($F_{(3, 72)} = 36.53$, $p < 0.001$) that the four regions differed in their sensitivity to different kinds of cognitive control. Specifically, the most dorsocaudal FEF preferred visuospatial control (hence, the label 'visual-sensitive), while the most ventrorostral IFG preferred semantic control (hence, the label 'semantic-sensitive'). Relative to the FEF and IFG that showed domain-specific preferences, the interposed regions (the MFG and IFS) did not exhibit any bias to either domain (hence, the label 'multiple-demand'). Asterisks *** $p < 0.001$ indicates a statistically significant positive bias (towards semantic control) or negative bias (towards visuospatial control). Non-significance (*n.s.*) indicates no bias.
